# Identification of Novel Gamma Radiation-Responsive Genes in the Thiosulfate-Oxidizing Bacterium *Limnobacter thiooxidans*

**DOI:** 10.64898/2026.05.23.727404

**Authors:** Tomoro Warashina, Asako Sato, Yuma Dotsuta, Toru Kitagaki, Takeshi Masuda, Haruo Ikeda, Masakazu Kataoka, Teppei Morita, Akio Kanai

**Affiliations:** Institute for Advanced Biosciences, Keio University, Tsuruoka, Japan; Systems Biology Program, Graduate School of Media and Governance, Keio University, Fujisawa, Japan; Japan Atomic Energy Agency, Tokai, Japan; Technology Research Association for Next-Generation Natural Products Chemistry, Tokyo, Japan; Department of Biomedical Engineering, Graduate School of Shinshu University, Nagano, Japan; Department of Infectious Diseases, Kyorin University School of Medicine, Mitaka, Japan; Faculty of Environment and Information Studies, Keio University, Fujisawa, Japan

**Keywords:** gamma irradiation, *Limnobacter thiooxidans*, proteome, transcriptome, stress response

## Abstract

Ionizing radiation induces DNA damage and oxidative stress; however, the genes and molecular mechanisms involved in bacterial stress responses have not been sufficiently identified. In this study, we used *Limnobacter thiooxidans* strain CS-K2, which is the closest relative to the bacteria detected in torus room water at the Fukushima Daiichi Nuclear Power Plant according to 16*S* rRNA gene sequences, and evaluated its response to γ-ray irradiation using integrated transcriptomic and proteomic analyses. We identified three previously uncharacterized genes (*LT3105*, *LT3115*, and *LT3126*) that were strongly induced at the mRNA and protein levels. These genes exhibited low basal expression but were markedly upregulated by γ-ray irradiation. Notably, *LT3126* encodes a protein containing VIT (vault protein inter-α-trypsin) and VWA (von Willebrand factor type A) domains and showed the strongest induction. Overexpression of *LT3126* increased survival after 500 Gy irradiation by approximately 200-fold compared with the control bacteria, demonstrating a direct contribution to survival under high-dose stress. Comparative genomic analysis showed that these genes are not widely conserved across bacteria but are unevenly distributed among specific lineages. Taken together, this study identified a novel set of γ-ray–responsive genes and demonstrated a functional role for *LT3126* in radiation resistance, providing new insights into molecular adaptation in radiation-associated environments.

**IMPORTANCE:** We identified a novel set of γ-ray–responsive genes (*LT3105*, *LT3115*, and *LT3126*) in the non-model bacterium *Limnobacter thiooxidans*. These genes are located in relatively close genomic proximity and are coordinately induced upon irradiation, suggesting a shared functional role in stress response. Overexpression of *LT3126* increased survival by approximately 200-fold after 500 Gy irradiation compared with the control bacteria, demonstrating a substantial contribution to survival under high-dose stress. These genes were also induced by heat shock and oxidative stress, indicating that their function extends beyond radiation-specific responses to broader environmental stress adaptation. Consistent with this, comparative genomic analysis showed that these genes are not widely conserved across bacteria but are unevenly distributed among specific lineages. Taken together, these findings highlight previously unrecognized molecular strategies that may support bacterial survival in radiation-associated environments.

---

Ionizing radiation damages DNA and proteins through direct molecular ionization and the generation of reactive oxygen species produced by water radiolysis (1). However, certain microorganisms that inhabit natural and artificial radiation environments can survive and proliferate under such conditions (2), making them important model systems for understanding the molecular basis of radiation resistance and oxidative stress responses (1). In recent years, extensive research into microbial communities in storage water containing radioactive elements and on metallic surfaces has revealed that a limited number of bacterial genera dominate, particularly in artificial closed water systems (3).

We previously analyzed the microbial communities in radioactive water environments using 16*S* rRNA amplicon sequencing, and commonly detected the thiosulfate-oxidizing bacterium *Limnobacter thiooxidans* (4). We also determined the complete genome sequence of *L. thiooxidans* strain CS-K2, a cultured isolate that is the closest relative to bacteria detected in radioactive water environments according to 16*S* rRNA gene sequence similarity (5). Notably, members of this species were independently detected in water storage systems in Japan and in spent nuclear fuel storage pools in France (6), suggesting the ability to adapt to radiation-associated environments. *L. thiooxidans* is an aerobic Gram-negative bacterium that utilizes thiosulfate as an electron donor and is widely distributed in both freshwater and marine environments (7). However, it is unclear how this organism responds to radiation stress and which molecular mechanisms support its survival in these conditions. Ionizing radiation not only damages DNA directly but also induces multiple intracellular responses, including oxidative stress and protein damage, through the generation of radical species via water radiolysis (8). Therefore, an integrative approach that captures these diverse cellular responses is required to understand the molecular basis of radiation resistance.

Previous studies of radiation-resistant microorganisms, including the Gram-positive bacterium *Deinococcus radiodurans* (9) and the thermophilic archaeon *Thermococcus gammatolerans* (10), have revealed key molecular mechanisms involved in radiation resistance, including genome multipartitioning, robust antioxidative systems, and highly efficient DNA repair machinery centered on RecA (11). However, most of these findings were based on a limited number of model organisms, and much remains unknown about how non-model microorganisms that inhabit diverse environments adapt to extreme conditions using previously uncharacterized functional molecules.

Therefore, in this study we investigated the responses to γ-ray irradiation of *L. thiooxidans* at the molecular level using integrated transcriptomic and proteomic analyses. We identified three previously uncharacterized genes (*LT3105*, *LT3115*, and *LT3126*) that responded strongly to radiation and oxidative stress. We describe their domain architectures, the radiation resistance conferred by *LT3126* overexpression, and their phylogenetic distribution based on comparative genomic analysis. These findings suggest that this organism adapts to radiation stress through strong induction of DNA repair-related pathways and changes in the expression of previously uncharacterized proteins.

## RESULTS AND DISCUSSION

### Transcriptomic and proteomic changes in *L. thiooxidans* exposed to γ-ray irradiation

We performed integrated transcriptomic and proteomic analyses to characterize the molecular response of *L. thiooxidans* CS-K2 exposed to γ-ray irradiation. Cobalt-60 (^60^Co) was used as the radiation source in three experimental conditions (Fig. 1A): non-irradiated (LR1; 0 Gy), 0.52 kGy (LR2), and 1.1 kGy (LR3). Irradiation was conducted for 11.8 days at dose rates of 1.82 and 3.9 Gy/h. Based on previous findings showing that this organism can survive at cumulative doses up to 700 Gy (4), we applied low-dose-rate irradiation over 11.8 days to approximate chronic exposure conditions, yielding cumulative doses of 500 Gy (sublethal) and 1100 Gy (high stress), while avoiding complete lethality. The gene locus tags, gene IDs, and functional annotations used in this study are summarized in Table S1. Following quality control (QC) of the sequencing and mass spectrometry data, the reads were mapped to the complete genome, resulting in the detection of 3,192 genes corresponding to all predicted open reading frames (ORFs) (Table S2A). Proteomic analysis identified 1,906–2,046 proteins, representing ∼60% of the predicted ORFs (Table S2B), indicating substantial coverage of mRNA and protein changes. We then compared the global transcriptomic and proteomic profiles. Principal component analysis (PCA) (12) (Fig. S1A and S1B) showed clear separation between the non-irradiated and irradiated samples, with the two irradiation conditions forming closely related clusters. This indicates that γ-ray irradiation was the primary driver of the variation in the global transcriptomic and proteomic profiles and that similar responses were induced at both doses. Consistently, the gene expression changes were highly correlated between the irradiation conditions (R^2^ = 0.89; Fig. S1C), and similar trends were observed in the proteome (Fig. S1B and S1D). Based on these results, the subsequent analyses focused on the 0.52 kGy condition as the lowest dose that elicits a robust response.

**FIG 1.**
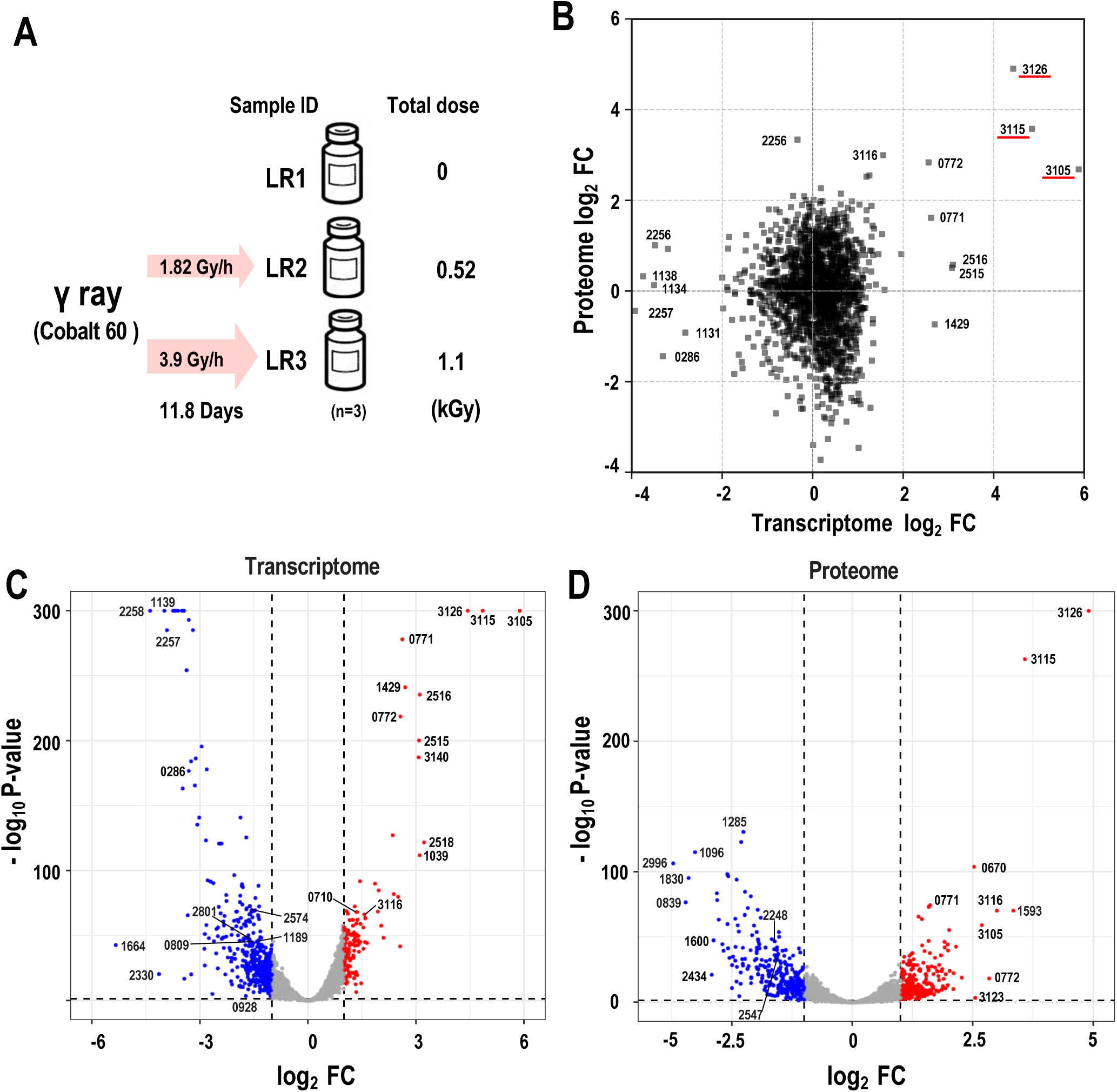
Transcriptomic and proteomic analyses of L. thiooxidans following γ-ray irradiation. (A) Experimental design. γ-ray irradiation using 60Co was performed under three conditions: no irradiation (LR1), 0.52 kGy (LR2), and 1.1 kGy (LR3). The dose rates were 1.82 and 3.9 Gy/h, with an irradiation period of 11.8 days. (B) Comparison of the log2FC at the transcriptomic (x-axis) and proteomic (y-axis) levels. (C) Volcano plot of RNA-seq. The x-axis shows the log2FC of gene expression in the 0.52 kGy-irradiated sample relative to the non-irradiated control, and the y-axis shows the −log10(p-value). Red dots represent significantly upregulated genes, blue dots represent significantly downregulated genes, and gray dots represent genes with no significant changes in expression. For representative genes, the numerical part of the gene ID is indicated (see Table S1). (D) Volcano plot based on proteomic analysis. The x-axis shows the log2FC of protein abundance in the 0.52 kGy-irradiated sample relative to the non-irradiated control and the y-axis shows the −log10(p-value). Color coding and numbering are as in (C).

We also compared transcriptomic and proteomic changes at 0.52 kGy and found a modest correlation between mRNA and protein fold changes (R² = 0.417; Fig. 1B). We then evaluated the global transcriptional response to γ-ray irradiation. In a volcano plot (Fig. 1C), 131 of 3,192 genes were significantly upregulated (log^2^ fold-change [log2Fc] ≥ 2.0, p < 0.05), whereas 393 genes were downregulated (log2FC ≤ −2.0, p < 0.05) relative to the non-irradiated condition. This pattern indicates the selective induction of a limited gene set, alongside broader repression. The upregulated genes (Table 1A) included DNA repair-related factors (13, 14) such as *RecA* (*LT0771*) and *RecX* (*LT0772*) (Fig. S2), which increased by 6.2- and 5.9-fold at the RNA level, respectively, confirming activation of the DNA damage response. Visualization of the log2FC data across genomic loci (Fig. 2A) showed that the induced genes were not uniformly distributed but were instead concentrated in specific regions. Notably, three previously uncharacterized genes (*LT3105*, *LT3115*, and *LT3126*) exhibited stronger induction than the known factors, with increases of 58.9-, 28.8-, and 21.6-fold, respectively, suggesting they have specific roles in the radiation response of *L. thiooxidans*.

**FIG 2.**
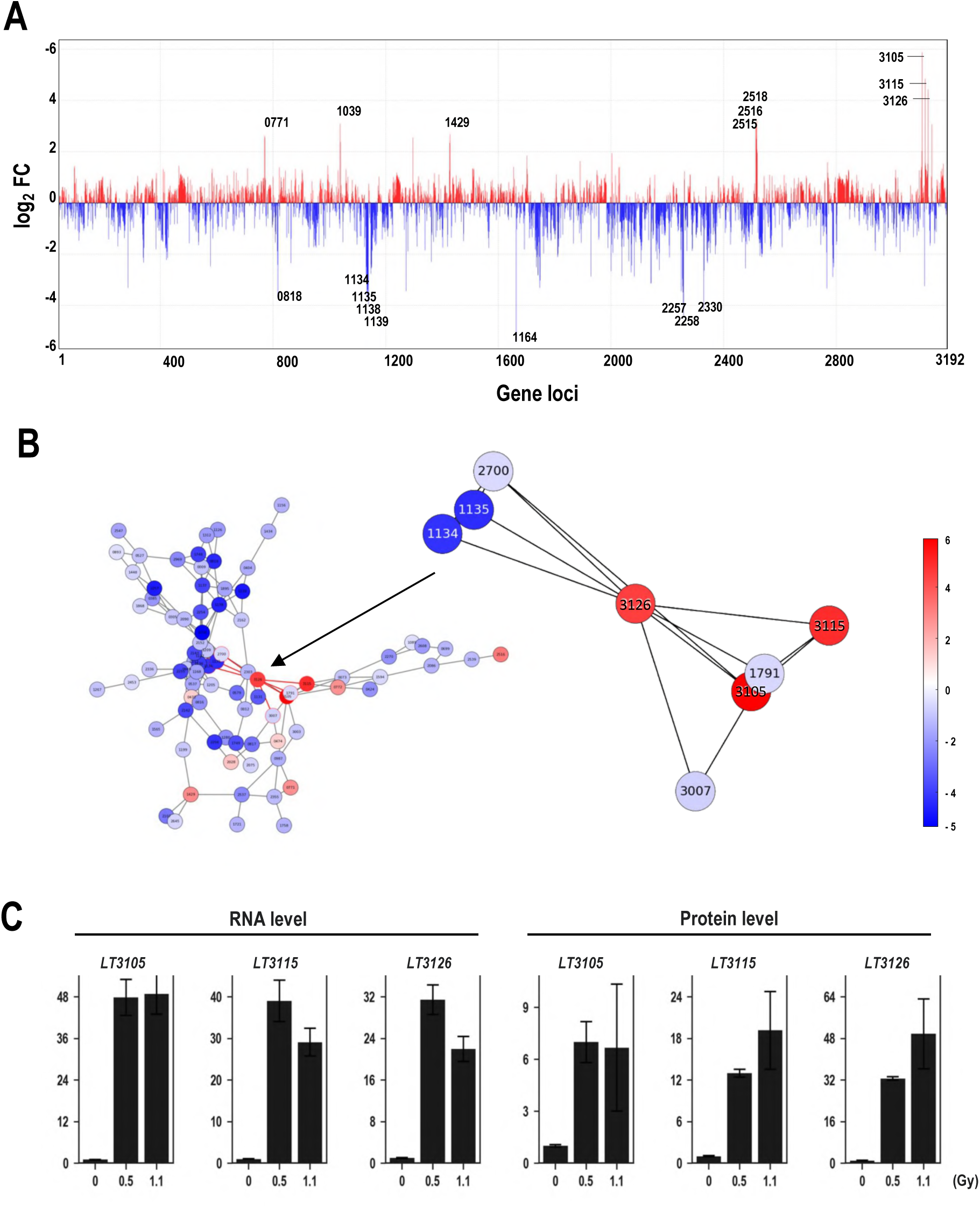
Transcriptomic alterations and gene network structure in *L. thiooxidans* following γ-ray irradiation. (A) Changes in gene expression along the *L. thiooxidans* genome after γ-ray irradiation. The *x*-axis represents gene loci (the locus numbers correspond to the gene IDs) and the *y*-axis represents the log2FC of expression in the 0.52 kGy-irradiated sample relative to the non-irradiated control. Red and blue colors indicate upregulated and downregulated genes, respectively. For representative genes, the numerical part of the gene ID is indicated. (B) Network analysis of DEGs and co-regulated gene clusters. Each node represents a gene, and the similarity of expression-change patterns is based on distance metrics. The node color indicates the direction of change in expression: red for upregulated and blue for downregulated genes. The color bar scale indicates the log2FC. The inset shows an enlarged view centered on gene *LT3126*, highlighting its eight directly connected nodes. (C) Changes in the mRNA and protein abundance of *LT3105*, *LT3115*, and *LT3126* after γ-ray irradiation. The *x*-axis indicates the irradiation dose (Gy) and the *y*-axis represents normalized mRNA and protein abundance (normalized counts). Error bars indicate one standard error.

**TABLE 1.**
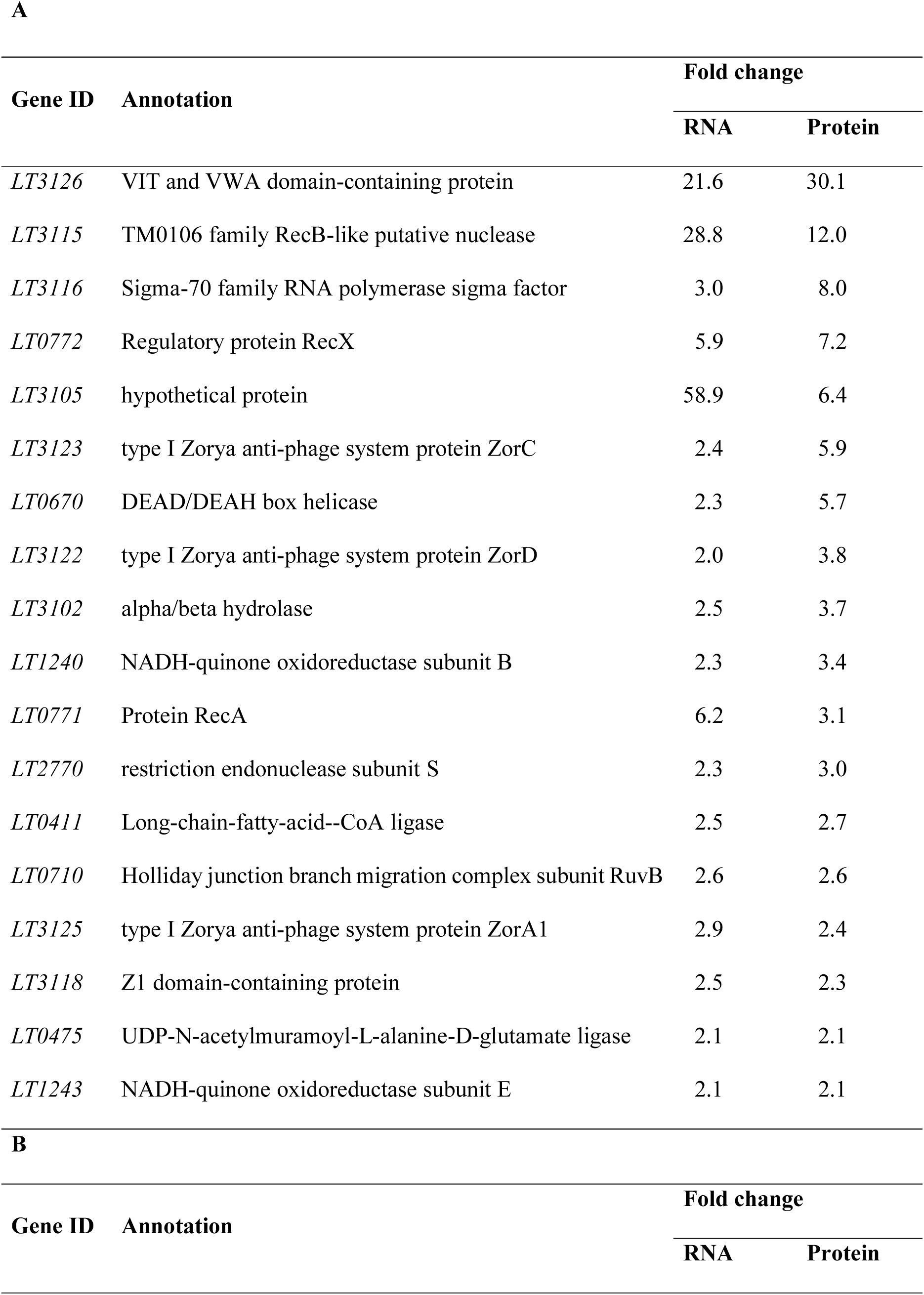
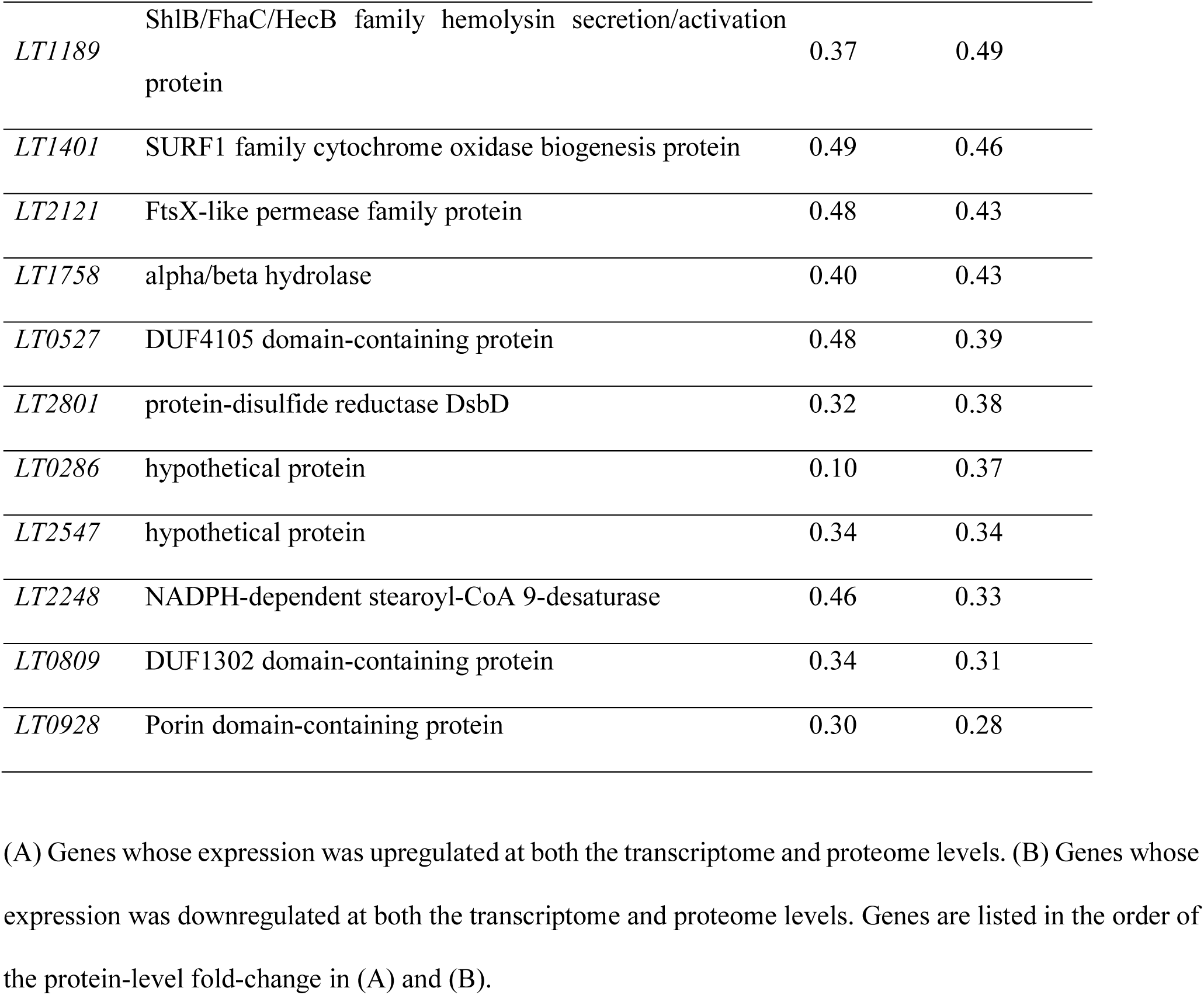
Genes showing significant changes in expression upon γ-ray irradiation in *L. thiooxidans*. (A) Genes whose expression was upregulated at both the transcriptome and proteome levels. (B) Genes whose expression was downregulated at both the transcriptome and proteome levels. Genes are listed in the order of the protein-level fold-change in (A) and (B).

To characterize the functional trends, we performed Clusters of Orthologous Genes (COG)-based (15) classification (Fig. S3). The upregulated genes were enriched in replication and repair (L), consistent with DNA damage response activation, as well as in translation (J) and energy production (C). In contrast, the downregulated genes were enriched in lipid transport and metabolism (I), and energy production and conversion (C), indicating the suppression of membrane and metabolic functions. These results suggest that the transcriptional response to γ-ray irradiation involves the activation of DNA repair pathways and broad repression of cellular metabolism, reflecting adaptive resource reallocation.

We also assessed the protein-level responses in a similar manner. The proteome analysis (Fig. 1D) revealed extensive changes in protein abundance, consistent with the transcriptome. Among ∼2,000 identified proteins, 219 were significantly increased and 256 were significantly decreased according to the log2FC (≥ 2.0 or ≤ −2.0, p < 0.05). The upregulated proteins (Table 1A; Fig. S2) included DNA repair factors, such as RecA (LT0771) (14), RecX (LT0772) (14), and RuvB (LT0710) (16), as well as the three previously uncharacterized proteins (LT3105, LT3115, and LT3126), which showed strong induction at the transcript level. In contrast, the downregulated proteins (Table 1B) were predominantly membrane-associated, including porin (LT0928) (17), NADPH-dependent stearoyl-CoA 9-desaturase (LT2248), and the redox protein DsbD (LT2801) (18). The COG analysis (Fig. S3) confirmed that the suppression of the membrane- and metabolism-related categories (M and C) observed at the transcript level was also reflected in the proteome. As noted above, the correlation between mRNA and protein fold changes was modest, suggesting that post-transcriptional regulation, including translational control and protein turnover, contributes to final protein abundance (19). Accordingly, the genes showing consistent and significant changes at both the transcriptomic and proteomic levels (Table 1) likely represent key components of the radiation response in *L. thiooxidans*.

### Gene expression network analysis and identification of novel γ-ray–responsive factors

To elucidate the regulatory architecture underlying the widespread expression changes, the distribution of differentially expressed genes (DEGs) across the genome and their relationships were analyzed. As shown in the genomic expression map (Fig. 2A; Fig. S4), the altered genes were distributed throughout the genome; however, in certain regions the upregulated genes tended to be in close proximity. In particular, local clustering of highly expressed genes was observed in several regions, including near *LT3126*. However, these genes are not continuously encoded and are instead considered to be expressed as independent transcriptional units.

To capture the relationships among these expression changes, we performed co-expression network analysis based on the similarity of gene expression patterns (Fig. 2B). In this analysis, each gene was represented as a node and the edges were defined based on the distance metrics of expression change patterns across the irradiation conditions. Consequently, the gene groups showing increased expression (red nodes) and those with decreased expression (blue nodes) tended to form distinct clusters. However, these subnetworks were not completely separated. Notably, in the enlarged network view (Fig. 2B), *LT3126* was connected to multiple upregulated genes, including *LT3115* and *LT3105*, as well as some downregulated genes, such as *LT1134* and *LT1135*. *LT1134* and *LT1135* encode homologs of *HscA* and *HscB,* respectively, which are involved in Fe-S cluster assembly. Because proteins containing the Fe-S cluster are highly sensitive to oxidative stress (20), the decreased expression of these genes may reflect disruption of the Fe-S cluster-related processes due to γ-ray irradiation. Although the co-expression network included both upregulated and downregulated genes, we focused on the strongly induced genes because they are more likely to represent active components of the stress response rather than the secondary effects of growth inhibition or metabolic suppression. The close proximity of these genes within the network indicates that they constitute a group of genes exhibiting similar expression responses under γ-ray irradiation. In general, genes that are closely positioned in co-expression networks have been reported to represent groups that undergo coordinated expression changes in response to a given environmental stimulus (19). The proximity of *LT3126*, *LT3115*, and *LT3105* in the network observed in this study suggests that these genes may be involved in a common stress response process under γ-ray irradiation.

To further support the importance of the three novel genes suggested by the network analysis, the response characteristics of the three novel genes (*LT3126*, *LT3105*, and *LT3115*) to different γ-ray doses were examined using the transcriptome and proteome data described above (Fig. 2C). These three genes exhibited extremely low expression at both the mRNA and protein levels under non-irradiated conditions but were clearly induced as the γ-ray dose increased. Compared with the known regulatory factor *recX* (RNA: 5.9-fold; protein: 7.2-fold), these genes exhibited higher induction at both the transcriptome and proteome levels, as exemplified by *LT3105* (RNA: 58.9-fold; protein: 6.4-fold), *LT3115* (RNA: 28.8-fold; protein: 12.0-fold), and *LT3126* (RNA: 21.6-fold; protein: 30.1-fold). The network positions and their pronounced increases in expression suggest that these novel genes may play important roles in the radiation response of *L. thiooxidans*.

### Domain structures and functional prediction of novel responsive proteins

To investigate and predict the domain structures of these novel gene products, we conducted structural prediction and domain analyses using AlphaFold2 (Fig. 3) (21). *LT3105* encodes a protein of 138 amino acids (14.3 kDa) with an unknown function. SignalP-6.0 analysis (22) predicted with high confidence the presence of an N-terminal Sec/SPII-type lipoprotein signal peptide (Fig. 3A). This suggests that the protein is likely to be a lipoprotein anchored to the cell membrane via lipid modification (23). In addition, the sequence comparisons with homologous proteins from other bacterial species confirmed that the N-terminal lipoprotein signal region is conserved (Fig. S5), suggesting that this structural feature may be evolutionarily conserved.

**FIG 3.**
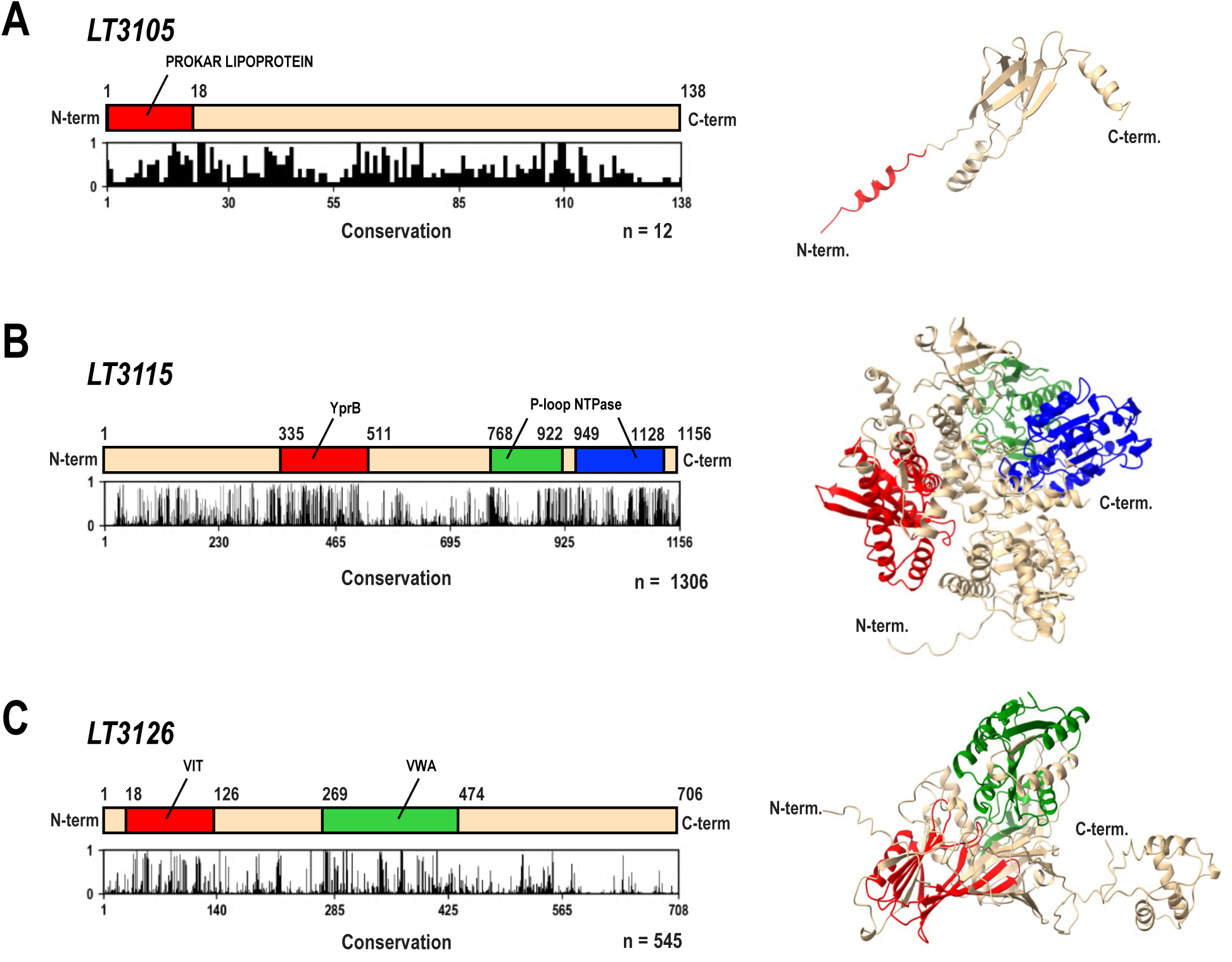
Predicted domain and structural features of LT3105, LT3115, and LT3126 proteins. The left column shows schematic representations of each protein, indicating the predicted domain positions and amino acid sequence conservation, while the right column presents the three-dimensional structures predicted by AlphaFold2 (60). The bar graphs below each protein schematic represent the conservation scores (61) of proteins with sequence similarity (see Methods), ranging from 0 to 1, where higher values indicate greater sequence conservation. (A) LT3105 protein. The PROKAR LIPOPROTEIN (Prokaryotic membrane lipoprotein lipid attachment site profile) domain is shown in red. (B) LT3115 protein. The YprB (ribonuclease H–like) domain is indicated in red. The P-loop NTPase (P-loop–containing nucleoside triphosphate hydrolase) domain corresponding to residues 706–945 is shown in green, and the region encompassing residues 959–1146 is shown in blue. (C) LT3126 protein. The VIT domain is shown in red and the VWA domain is shown in green.

Among the analyzed proteins, *LT3115* exhibited the most prominent multidomain structure. It encodes a protein of 1,156 amino acids (130.9 kDa) with an unknown function, and contains a helicase C-terminal domain (YprB [ribonuclease H–like]) (24) at the N-terminus and two P-loop_NTPase (P-loop–containing nucleoside triphosphate hydrolase) domains (25) at the C-terminus (Fig. 3B). Structural comparison analysis (Fig. S6) revealed that LT3115 shows high structural similarity to a RecB-like nuclease derived from *Mycobacterium leprae*, a bacterium known as the causative agent of leprosy, as well as to the eukaryotic DNA repair-associated DNA2/NAM7 helicase. Furthermore, the alignment analysis of homologous proteins (Fig. S7) and detailed domain analysis confirmed that the Walker A and Walker B motifs (26), which are essential for ATP hydrolysis, are highly conserved (Fig. S7). *RecB* nuclease is a central enzyme that functions as part of the *RecBCD* complex to process DNA double-strand break ends and initiate homologous recombination repair (27). The presence of this domain in *LT3115* suggests that this protein may possess enzymatic functions involved in the recognition and processing of damaged DNA.

*LT3126*, which showed the strongest induction, encodes a protein of 708 amino acids (77.5 kDa) with an unknown function. It was predicted to contain VIT (vault protein inter-α-trypsin) and VWA (von Willebrand factor type A) domains (Fig. 3C; Fig. S8). The VIT domain is a structural domain involved in protein–protein interactions and the regulation of complex functions (28). The VWA domain is widely conserved in cell adhesion factors such as integrins and collagen, and is known to mediate metal ion-dependent protein–protein interactions (28). The alignment analysis (Fig. S9) revealed the conservation of residues characteristic of the MIDAS (metal-ion-dependent adhesion site) motif, which is important for interactions, within the VWA domain of *LT3126*. Furthermore, the structural prediction indicated that the region encompassing amino acid residues 2–464, which contained the predicted domains, adopts a structure similar to that of uncharacterized proteins from *Variovorax paradoxus* (bacterium) and *Rotaria sordida* (rotifer) (Fig. S8). Although the functions of these proteins remain unknown, their conserved structural framework suggests they share common molecular functions. Based on these findings, *LT3126* is suggested to function not as an enzyme with independent catalytic activity but rather as a factor that acts via protein–protein interactions, and potentially participates in the formation and stability regulation of protein complexes within the cytoplasm.

### Overexpression of the novel VWA domain-containing gene *LT3126* confers γ-ray resistance

As described above, the omics expression of *LT3126* was markedly increased, and it was predicted to contain a protein–protein interaction domain (VWA) based on the predicted structure. However, the increased omics expression represents only a correlation with irradiation and does not necessarily indicate a direct causal role in the acquisition of resistance. Therefore, to directly verify whether the intracellular abundance of *LT3126* is a determinant in radiation resistance, we performed functional analysis using an overexpression system.

During the construction of the expression system, we attempted to introduce genes by transformation, but no transformants were obtained. Therefore, a system enabling stable and high-level expression of *LT3126* was established by conjugation. The broad-host-range plasmid pRSF1010CT was used as the vector. The plasmid, pLT3126OE, was constructed by incorporating the upstream region of the ribosomal protein L10 (*rplJ*) gene of *L. thiooxidans*, which is expected to confer constitutive high expression, as the promoter (Fig. 4A and 4B; Fig. S10). To confirm the expression level of *LT3126* in the constructed strains, quantitative RT-PCR was performed using total RNA extracted from the strain harboring pLT3126OE (overexpression strain) and the strain harboring the empty vector (control strain). This analysis revealed that the mRNA level of *LT3126* was significantly increased, by approximately 5-fold, in the overexpression strain compared with the control strain (Fig. 4C). The growth rate of the overexpression strain (Fig. 4D) was not significantly different (p < 0.05) to that of the control strain within the optical density (OD) range of 0.2–1.2 during the exponential growth phase. In contrast, in the late phase of culture, the OD decreased in the control strain (pRSF1010CT), but this decrease was relatively suppressed in the overexpression strain. However, under the experimental conditions used in this study, the factors underlying this difference could not be identified. These observations suggest that even when LT3126 protein accumulates excessively within the cell, it does not interfere with normal cell division or metabolic processes. This indicates that LT3126 might specifically function under stress conditions.

**FIG 4.**
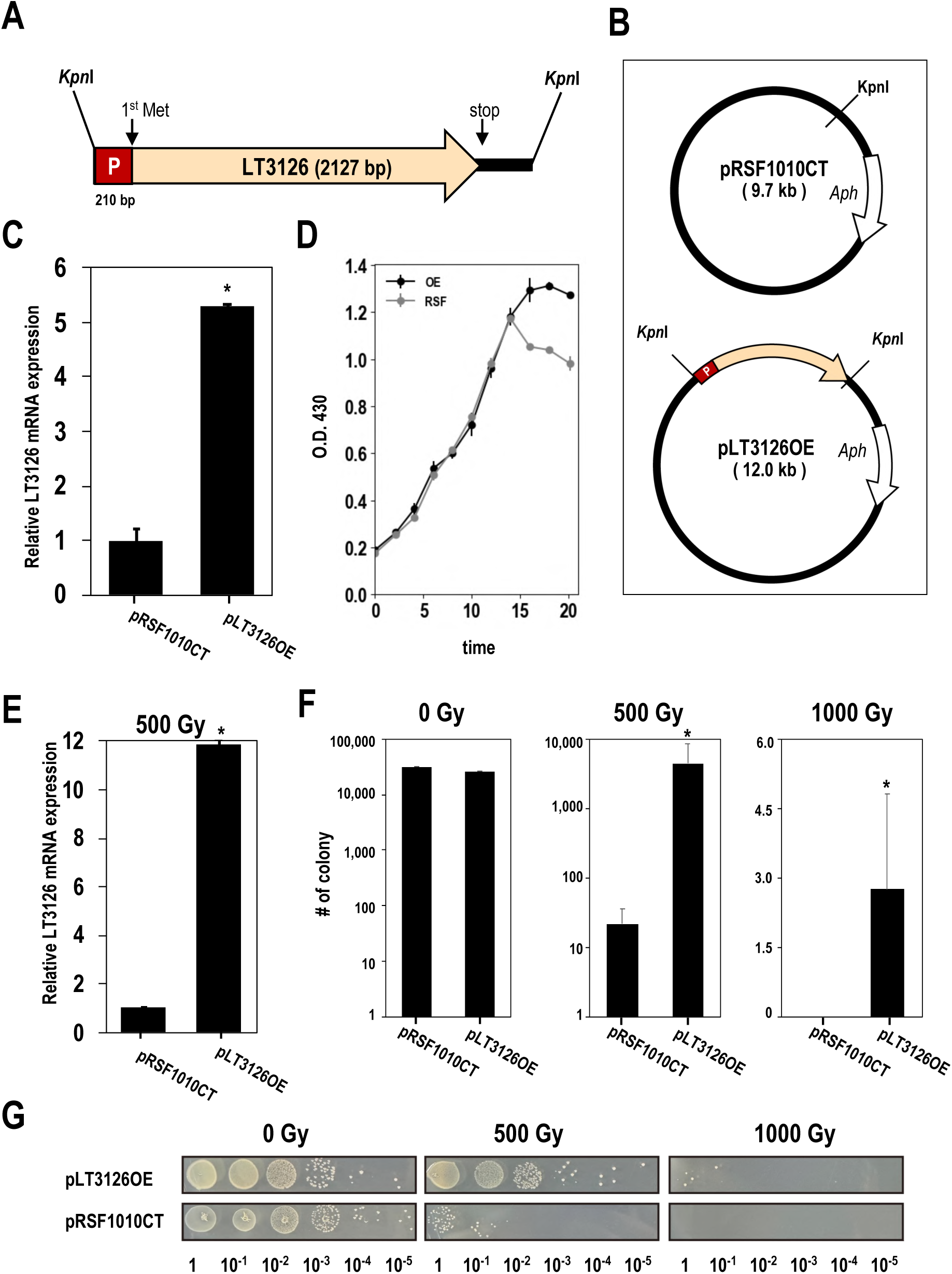
Overexpression of the *LT3126* gene enhances gamma-ray resistance in *L. thiooxidans*. (A) Schematic representation of construction of the *LT3126* overexpression cassette. The promoter region (P; shown in brown) of the ribosomal protein L10 gene derived from *L. thiooxidans* and the *LT3126* gene (beige) were inserted via a *Kpn*I restriction site. The start codon (first Met) and stop codon are indicated. (B) Schematic representations of the vector plasmid pRSF1010CT (9.7 kb) and the *LT3126* overexpression plasmid pLT3126OE (12.0 kb). *Aph* indicates the kanamycin resistance gene (aminoglycoside 3′-phosphotransferase). The *Kpn*I sites are indicated. (C) Expression profile of *LT3126* mRNA under non-irradiated conditions in *L. thiooxidans* introduced with pRSF1010CT or pLT3126OE. (D) Comparison of the growth rates of the control strain and the overexpression strain under non-irradiated conditions. Growth curves of the strains introduced with pRSF1010CT (gray) or pLT3126OE (black) in liquid culture are shown. (E) Expression profile of *LT3126* mRNA following 500 Gy irradiation in *L. thiooxidans* introduced with either pRSF1010CT or pLT3126OE. (F) Quantitative analysis of the colony formation assay shown in (G). Colony-forming units were counted under each irradiation condition. The *y*-axis indicates the number of colonies; data for 0 and 500 Gy are plotted on a logarithmic scale, whereas data for 1000 Gy are plotted on a linear scale. Error bars represent the standard deviation (SD). (G) Survival of *L. thiooxidans* introduced with pRSF1010CT or pLT3126OE following γ-ray irradiation. After exposure to the indicated doses (0, 500, and 1000 Gy), the cultures were serially diluted (10^0^–10⁻^5^) and plated on agar medium. Representative colony morphologies are shown. Asterisks (*) indicate statistically significant difference compared with the pRSF1010CT strain (p < 0.05; two-tailed Student’s *t* test).

To examine whether the expression level of *LT3126* affects radiation resistance, cultures in the exponential growth phase were dispensed into vials and subjected to γ-ray irradiation for approximately 10 days. After irradiation, the surviving colonies were counted, and mRNA levels were analyzed. Quantitative RT-PCR analysis confirmed that the expression level of LT3126 was significantly greater in the overexpression strain compared with the control strain following 500 Gy irradiation (Fig. 4E). Under non-irradiated conditions (0 Gy), there was no significant difference in the colony counts, and the difference between the two strains was minimal (Figs. 4F and 4G). In contrast, following γ-ray irradiation, the survival clearly differed between the two strains. After 500 Gy irradiation, the mean colony count was 4.4 × 10^3^/mL for the control strain compared with 9.0 × 10^5^/mL for the overexpression strain, representing a significant increase in survival by approximately 200-fold (Figs. 4F and 4G). Furthermore, under 1000 Gy irradiation, which is close to the survival limit of this organism, no surviving colonies were detected in the control strain, whereas 5.5 × 10^2^/mL colonies were counted for the overexpression strain (Figs. 4F and 4G). These results indicate that increased abundance of *LT3126* contributes to survival under extreme conditions.

To evaluate the specificity of the stress response of *LT3126*, focusing on the early stress response, the changes in mRNA expression over short time periods under heat shock and oxidative stress conditions were analyzed by RT-PCR (Fig. 5). Under oxidative stress, *LT3126* expression increased by 2.09-fold at 15 min and 2.31-fold at 30 min. Stronger induction was observed under heat shock, with a 3.65-fold increase at 15 min and a 3.27-fold increase at 30 min. Similarly, *LT3115* exhibited marked induction, with increases of 13.20-fold (15 min) and 14.80-fold (30 min) under oxidative stress, and 2.29-fold (15 min) and 3.36-fold (30 min) under heat shock, indicating that both genes exhibit similar responsiveness depending on the type of stress. In contrast, *LT3105* did not show a significant response to heat shock, with specific induction under oxidative stress (Fig. 5B). The control genes used as stress indicators, hsp70 (heat-responsive chaperone) (29) and recX (a regulator of RecA) (14), were induced in response to both types of stress, confirming that these environmental stresses were appropriate in this experimental system. These results strongly suggest that the analyzed factors, including *LT3126*, retain a certain level of responsiveness not only to radiation but also to general environmental stresses, such as heat and oxidative stress.

**FIG 5.**
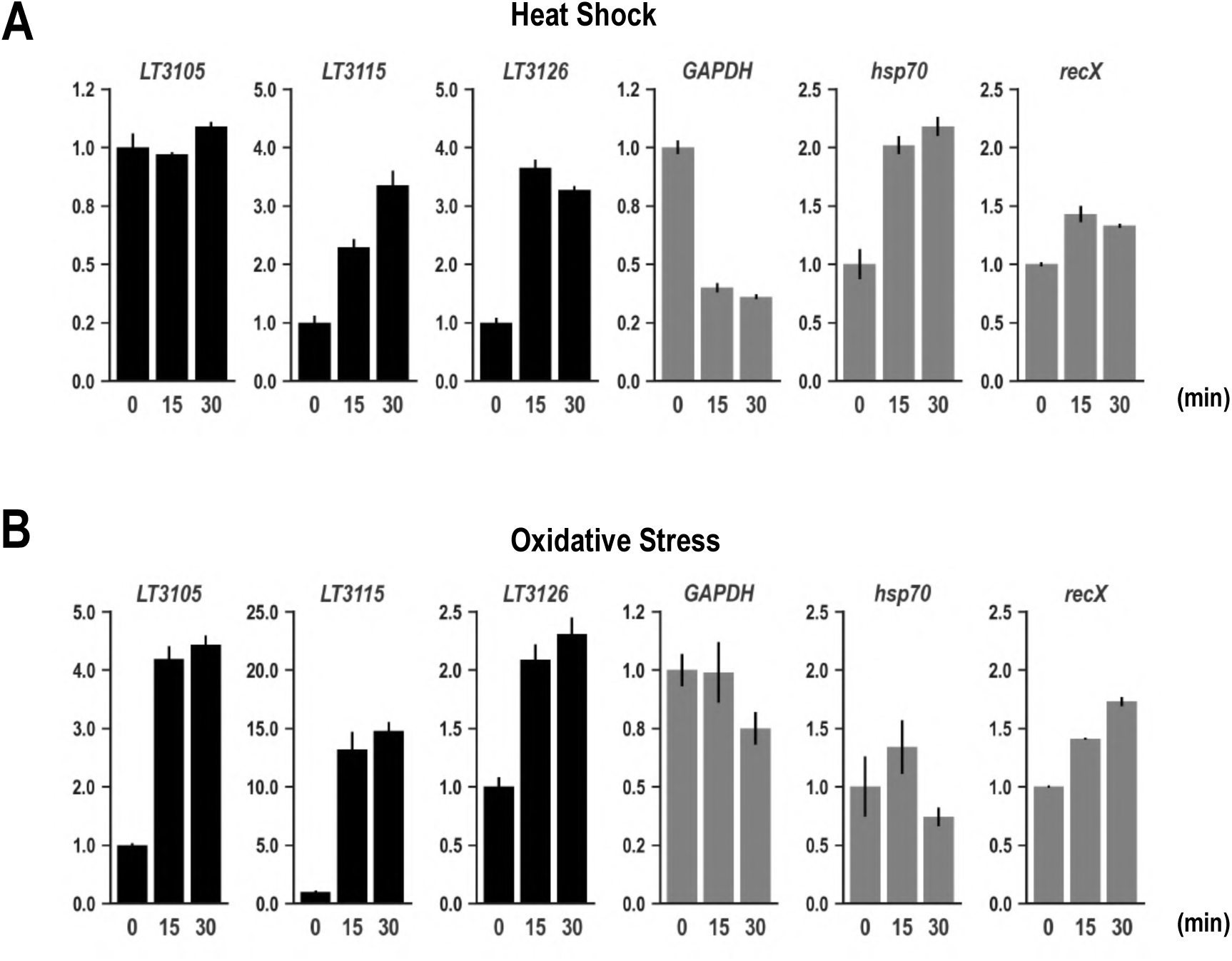
Changes in the expression of three *L. thiooxidans* genes under heat shock and oxidative stress conditions. (A) mRNA expression profiles of *LT3105*, *LT3115*, and *LT3126* under heat shock conditions. (B) mRNA expression profiles of *LT3105*, *LT3115*, and *LT3126* under oxidative stress conditions. *GAPDH*, *hsp70*, and *recX* were used as reference controls. The bar graphs show the mRNA expression levels at 0, 15, and 30 min. The bars for *LT3105*, *LT3115*, and *LT3126* are shown in black, while those for the reference controls are shown in blue. The *y*-axis shows the relative mRNA expression levels, and error bars indicate the standard error.

On the other hand, attempts to construct a gene knockout strain of *LT3126* did not yield viable transformants. This may be explained by the possibility that *LT3126* is an essential gene required for growth under normal conditions as well as technical limitations, such as low efficiency of conjugation and homologous recombination, which prevented sufficient screening size for identifying gene disruption mutants. Therefore, further optimization of the experimental system is required to clarify the physiological function of *LT3126*.

### Evolutionary distribution of the *LT3105*, *LT3115*, and *LT3126* gene set

The analyses described above suggest that, while *LT3105*, *LT3115*, and *LT3126* are functionally differentiated, these genes may form a novel stress response system that is coordinately induced under γ-ray irradiation (Fig. 2C) and oxidative stress (Fig. 5B). To elucidate the evolutionary background of this system, we performed a large-scale comparative genomic analysis using 5,927 bacterial reference genomes registered in the NCBI RefSeq database (30) (Table S3). First, we examined the broad distribution of these three genes across bacteria (Fig. S11), and found that none of them are universally conserved core genes, but are instead unevenly distributed among specific taxonomic groups. Mapping the genes onto the phylogenetic tree showed that they are not uniformly distributed across the bacterial domain; instead, they are sporadically detected in certain clades within Proteobacteria and Actinobacteria, but completely absent in many other lineages. Furthermore, the presence or absence of these genes was mixed, even among closely related taxa, and no continuous phylogenetic conservation was observed.

Next, the analysis of the co-occurrence relationships of these genes at the genus level revealed marked differences in their distributions (Fig. S12). Notably, *LT3115* was independently detected in 968 genera and exhibited the widest distribution among the three genes, whereas *LT3126* was found in 344 genera and *LT3105* in only 12 genera, indicating a pronounced asymmetry in their distribution breadth. The co-occurrence of two genes was limited; *LT3115* and *LT3126* co-occurred in 124 genera, whereas combinations including *LT3105* were extremely rare. Only one genus (*Limnobacter*) possessed all three genes simultaneously. These findings suggest that these genes are not consistently maintained as a single functional module but may have independent evolutionary histories and were only reassembled in specific lineages.

To provide a contrast with this broad-scale rarity, local conservation within the genus *Limnobacter* (38 strains) was examined in detail (Fig. S13). This analysis revealed that *LT3105* was relatively widely conserved within the genus, being present in the major clade containing *L. thiooxidans* (27 strains) and lower branches, such as *L. profundi*. In contrast, *LT3115* and *LT3126* were specifically conserved in the upper clade of the phylogenetic tree, which includes *L. thiooxidans* and the closely related species *Limnobacter* sp. E19-10.1d, and were consistently absent in lower clades, such as *L. profundi*. This distribution suggests an evolutionary process in which *LT3105*, originally present in the ancestor of the genus *Limnobacter*, was accompanied by the acquisition of the *LT3115* and *LT3126* module in specific lineages, possibly through horizontal gene transfer (31). This pattern further indicates that these genes are not uniformly conserved across the bacterial domain but reached their present distribution through lineage-specific evolutionary processes. Since members of the genus *Limnobacter* are widely distributed in aquatic environments, including the deep sea, volcanic sediment, and environments contaminated with radioactive materials (4, 32, 33), it is possible that they acquired unique defense mechanisms during adaptation to such environmental stresses.

## Conclusions

We identified a novel set of γ-ray–responsive genes (*LT3105*, *LT3115*, and *LT3126*) in *Limnobacter thiooxidans* strain CS-K2 and characterized their functional and evolutionary features. The integrated transcriptomic and proteomic analyses showed that these genes exhibit low basal expression but are strongly induced at both the mRNA and protein levels upon exposure to irradiation and are coordinately regulated in association with DNA repair-related genes. The functional analysis demonstrated that *LT3126* overexpression significantly enhanced survival under γ-ray irradiation, with an approximately 200-fold increase in colony-forming units at 500 Gy, indicating a direct contribution to survival under high-dose stress. In addition to radiation, these genes also responded to heat shock and oxidative stress, suggesting a broader role in adapting to environmental stress. The comparative genomic analysis further revealed that these genes are not widely conserved across bacteria but are instead unevenly distributed among specific lineages, thereby supporting lineage-specific acquisition and retention. In summary, this study has identified previously unrecognized γ-ray–responsive genes and provides new insights into the molecular strategies that support bacterial survival in radiation-associated and oxidative stress environments.

## MATERIALS AND METHODS

### Cultivation of *Limnobacter thiooxidans* and ionizing radiation resistance assay

We used *Limnobacter thiooxidans* strain CS-K2, which was obtained from the Deutsche Sammlung von Mikroorganismen und Zellkulturen (DSMZ; Braunschweig, Germany). Cells were cultivated in Marine Broth 2216 (Difco, BD, Franklin Lakes, NJ, USA) at 30 °C with shaking at 150 revolutions per minute (rpm). After 20 h of incubation, the culture was diluted 100-fold with fresh medium, and 60 mL aliquots were dispensed into sealable sterile glass vials (0501-09; Maruemu Corporation, Osaka, Japan). The diluted cultures were statically incubated for ∼48 h until the OD reached 0.4–0.5. Ionizing radiation resistance assays using a ^60^Co γ-ray source were conducted at the Takasaki Advanced Radiation Research Institute, Takasaki, Japan. Three irradiation conditions were applied: a high dose rate (3.9 Gy/h), a low dose rate (1.82 Gy/h), and a non-irradiated control (0 Gy/h). For each condition, three independent biological replicates were prepared (*n* = 3). Irradiation was performed continuously for 11.8 days, during which time the vials were maintained under static incubation at room temperature. After irradiation, the cells were harvested by centrifugation (8,000 rpm, 10 min, 4 °C) for subsequent analyses.

### Transcriptomic analysis

#### RNA preparation

The harvested cells were resuspended in TE–lysozyme buffer (1 mL per 10 mL of culture; 13 mg/mL lysozyme in TE buffer [10 mM Tris-HCl, 1 mM EDTA, pH 8.0]). The suspension was kept on ice, briefly vortexed to homogenize, and then incubated at room temperature for 5 min. Buffer RLT (Qiagen, Hamburg, Germany) supplemented with β-mercaptoethanol was added, and the cells were disrupted on ice using a Polytron PT2100 homogenizer (Kinematica, Lucerne, Switzerland) with three bursts of 10–20 s each. An equal volume of phenol:chloroform:isoamyl alcohol (25:24:1) was added to the lysate, mixed thoroughly, and centrifuged (12,000 rpm, 4 °C, 10 min) to collect the aqueous phase. An equal volume of chloroform was added, followed by centrifugation under the same conditions to remove residual phenol. The aqueous phase was transferred to a new tube, mixed with 1/10 volume of 3 M sodium acetate (pH 5.2) and 2.5 volumes of 100% ethanol, and incubated at −20 °C for 60 min to precipitate total RNA. RNA was pelleted by centrifugation (14,000 rpm, 4 °C, 20 min), washed twice with 20 mL of ice-cold 75% ethanol, and re-centrifuged (14,000 rpm, 4 °C, 5 min). After removing the supernatant, the pellet was briefly air-dried and dissolved in RNase-free water. The RNA solution was treated with DNase I (Takara Bio, Shiga, Japan), purified with phenol:chloroform:isoamyl alcohol (25:24:1), and centrifuged (15,000 rpm, 4 °C, 10 min). The final RNA pellet was resuspended in 100 µL of RNase-free water and stored at −80 °C.

#### cDNA library preparation and sequencing

RNA-seq was outsourced to GeneBay Inc. (Yokohama, Japan). The cDNA library was prepared as follows. rRNA was removed from total RNA, after which the RNA was fragmented for cDNA synthesis. Strand specificity was introduced by incorporating dUTP during second-strand cDNA synthesis. The resulting double-stranded cDNA underwent end repair, adapter ligation, enzymatic digestion, size selection, and PCR amplification to generate the sequencing libraries. The final libraries were sequenced on a NovaSeq 6000 platform (Illumina, San Diego, CA, USA) to obtain paired-end 150 bp reads (2 × 150 bp).

#### QC and expression analysis

The raw reads were processed using BBDuk (BBTools v38.90) (34) to trim low-quality bases and adapter sequences, and the remaining low-quality reads were removed using fqtrim (v0.9.4) (35). Default parameters were used for both tools. Reads with a mean quality score of <20 or a length of <50 bp were discarded. Clean reads were mapped to the *L. thiooxidans* CS-K2 reference genome (∼3.54 Mbp) using Bowtie2 (v2.4.5, --sensitive mode) (36). Gene-level read counts were obtained using featureCounts (v2.0.1) (37). The count read data were normalized, and statistical analyses of differential expression between conditions were performed using DESeq2 (v1.38.0, R v4.2.2) (38). DEGs were identified using a false discovery rate threshold of <0.05 after Benjamini–Hochberg correction. Functional annotation was performed using eggNOG-mapper (v2.1.9, database v5.0; web-based) (39) to assign Gene Ontology (GO) terms. GO enrichment analysis was used to identify enriched metabolic categories. Additionally, the COG database (15) was used to map genes to relevant metabolic pathways.

### Proteome analysis

Proteins were extracted using phase-transfer surfactant (PTS) buffer containing 12 mM sodium deoxycholate (SDC), 12 mM sodium lauroylsarcosinate (SLS), and 100 mM Tris-HCl (pH 9.0), and then quantified using a bicinchoninic acid assay kit. We used 10 μg of protein for digestion. Proteins were reduced and alkylated with 10 mM dithiothreitol and 50 mM chloroacetamide, respectively. Protein solutions were diluted 3-fold with 50 mM ammonium bicarbonate. Proteins were digested with Lys-C for 3 h followed by trypsin overnight at 37 °C. SDC and SLS were removed from the peptide solution using the PTS method (40, 41). Briefly, an equal volume of ethyl acetate was added to the peptide solution, and the mixture was acidified with trifluoroacetic acid (TFA). The mixture was agitated for 2 min and centrifuged to separate the organic and aqueous phases. The organic phase containing SDC and SLS was removed and dried. The dried peptides were resuspended in 5% acetonitrile containing 0.5% TFA and purified using StageTip (42). Nanoscale liquid chromatography–tandem mass spectrometry was performed using a mass spectrometer (Orbitrap Exploris 480; Thermo Fisher Scientific, Waltham, MA, USA) equipped with an ultrahigh-performance liquid chromatography system (Vanquish Neo; Thermo Fisher Scientific). The flow rate was 300 nL/min and the injection volume was 5 μL. An analytical column (Nikkyo Technos, Tokyo, Japan; inner diameter 75 μm, length 12 cm, packed with 3 μm C18 beads) was used. Data were acquired in the data independent mode. The raw data were converted to the mzML format using MSConvertGUI and the results files were analyzed using DIA-NN 1.8.1 with reference to the UniProt human proteome database.

### Stress-response assays

#### Heat shock treatment

*L. thiooxidans* CS-K2 was cultivated in Marine Broth 2216 at 30 °C with shaking at 150 rpm. Cells harvested after 22 h of incubation, corresponding to the exponential growth phase, were used for the heat shock experiments. Aliquots of 10 mL culture were transferred to a water bath preheated to 42 °C and subjected to heat shock for 0 min (control), 15 min, or 30 min. Three independent biological replicates were prepared for each condition. Following treatment, the cultures were immediately placed on ice and cells were collected by centrifugation (10,000 rpm, 4 °C, 3 min). Cells were harvested immediately after each treatment for RNA extraction.

#### Oxidative stress treatment

Using 10 mL aliquots of the same exponential-phase cultures as used for the heat shock experiments, oxidative stress was induced by adding hydrogen peroxide to a final concentration of 0.5 mM for 0 min (control), 15 min, or 30 min, with three independent biological replicates prepared for each condition. After treatment, the cultures were immediately placed on ice and cells were harvested by centrifugation (10,000 rpm, 4 °C, 3 min) for RNA extraction.

#### RT-PCR

cDNA was synthesized from 3 µg of total RNA using ReverTra Ace qPCR RT Master Mix (Toyobo, Osaka, Japan) with the manufacturer’s primer mix containing random primers and oligo(dT) primers. The resulting cDNA was used as the template for PCR amplification of the target genes (*LT3105*, *LT3115*, *LT3126*, *hsp70*, and *recX*) and the internal reference genes (*16S rRNA* and *GAPDH*). *GAPDH* encodes glyceraldehyde-3-phosphate dehydrogenase, *hsp70* encodes a heat-responsive chaperone, and *recX* encodes a regulator of RecA-mediated DNA repair. The primer sequences are listed in Table S5. PCR was performed using KOD FX Neo (Toyobo) under the following cycling conditions: initial denaturation at 94 °C for 2 min; 30 cycles of 98 °C for 30 s, 60 °C for 30 s, and 68 °C for 30 s; and final extension at 68 °C for 30 min.

#### Gel electrophoresis and quantification of band intensity

The PCR products were separated on 2% agarose gels and stained with ethidium bromide. Gels were imaged using a ChemiDoc XRS+ imaging system (Bio-Rad, Hercules, CA, USA) and band intensities were quantified using Image Lab software (Bio-Rad). For each gene, the band intensity was normalized to that of the internal reference gene (16*S* rRNA) and the relative expression levels were calculated by setting the control sample to a value of 1. All quantitative measurements were conducted using three independent biological replicates.

### Bioinformatics analysis

#### Statistical analysis and visualization of the omics data

For the transcriptomic and proteomic datasets, normalized expression values were used to assess the relationships among samples by PCA (12). To evaluate correlations in expression changes (log2FC) between the irradiation doses (0.5 vs. 1.1 kGy) and between the omics layers (transcriptome vs. proteome), Pearson’s correlation coefficients were calculated using the Python SciPy library (43). Volcano and MA plots (M, log2 fold change; A, log2 mean abundance) were generated to visualize the DEGs and differentially expressed proteins (DEPs). These analyses and visualizations were performed primarily using R and associated packages, such as DESeq2 (38) and ggplot2 (44).

#### Gene distribution and phylogenetic analysis

The phylogenetic distributions of the target genes were analyzed using 5,927 complete bacterial reference genomes available in the NCBI RefSeq database (30). The amino acid sequences encoded by *LT3105*, *LT3115*, and *LT3126* from *L. thiooxidans* CS-K2 were used as query sequences to identify <u>proteins</u> with high sequence similarity using BLASTp (45) with a threshold E-value of <1.0 × 10⁻^10^. When multiple hits were obtained within a single genome, the hit with the highest bit score was retained as the representative protein with the highest sequence similarity. The phylogenetic relationships among the 5,927 genomes were inferred using phyloT (46) based on NCBI Taxonomy. The presence/absence of each gene was mapped onto the phylogenetic tree using iTOL (47). For phylogenetic reconstruction within the genus *Limnobacter* (38 strains), GToTree (48) was used based on the core genome (*n* = 325). The conservation of orthologs in each strain was assessed by calculating the BLAST Score Ratio using LS-BSR (49).

#### Sequence and structural analyses

Conserved protein domains were predicted using InterProScan (50) against the InterPro database. When multiple domains or domain names were returned, the most representative domain was selected. Multiple sequence alignments of proteins with sequence similarity were generated using MAFFT (version 7) (51) and visualized using the ClustalX color scheme (52). The three-dimensional structures of the *L. thiooxidans* proteins (LT3105, LT3115, and LT3126) were predicted using ColabFold (53). Structural similarity searches of the predicted models were conducted against the AlphaFold Protein Structure Database using Foldseek (54). Structural superposition and visualization were carried out using UCSF ChimeraX (55). The statistical significance of differences in alignment lengths between bacterial and eukaryotic sequences was evaluated using Student’s *t* test.

### Construction of the *LT3126* overexpression strain

The *LT3126* overexpression strain of *L. thiooxidans* was constructed using the broad-host-range, self-replicating plasmid RSF1010CT. This plasmid is a derivative of pJRD215 (56) in which the *str* genes were replaced with the *cat* gene from pC194 (57) to confer chloramphenicol resistance, and further supplemented with the *oriT* region from pUB307 (58). To facilitate efficient plasmid transfer, the conjugative donor strain *E. coli* OMK64 (Δ(*mcrC-hsdSMR*), Δ(*srl-recA*), Δ*dcm*, Δ*dapA*, *rrnB*, Δ*lacZ4787 HsdR514* Δ(*araBAD*)*567* Δ(*rhaBAD*)*568 rph-1*) was engineered from its parental strain, BW25113, using a multistep lambda Red recombination system (59). OMK64 carries a pUB307 derivative in which the *aph* gene was precisely deleted via the same lambda Red system. A synthetic DNA fragment was generated by fusing a 210-bp promoter region upstream of the translation start site of the *L. thiooxidans* 50*S* ribosomal protein L10 gene (*LT2840*), which was identified in this study as a strong promoter, to the 2,127-bp coding sequence of *LT3126*. The fragment was commercially synthesized (Eurofins Genomics, Tokyo, Japan) with the addition of *Kpn*I restriction sites at both ends. The synthesized fragment was cloned into the *Kpn*I site of RSF1010CT to generate the overexpression plasmid pLT3126OE (Fig. 4A). For plasmid transfer, pLT3126OE was introduced into *E. coli* OMK64 harboring pUB307, which served as the donor strain. Donor cells and *L. thiooxidans* CS-K2 recipient cells were grown to the exponential phase, mixed at a 3:1 ratio, and incubated in Marine Broth at 30 °C for 5 h to allow conjugation. After incubation, the cells were resuspended, serially diluted, and plated onto R2A agar containing kanamycin (25 µg/mL). Transconjugants carrying pLT3126OE were selected and used for subsequent analyses.

## ACKNOWLEDGMENTS

We thank all members of the RNA Group at the Institute for Advanced Biosciences, Keio University, Japan, for insightful discussions. This work was supported in part by a KAKENHI Grant-in-Aid awarded to Japan for the Society for the Promotion of Science (JSPS) Fellows (22J22913) (to TW), and research funds from the Yamagata Prefectural Government and Tsuruoka City, Japan. The funding bodies played no role in the study design, data collection or analysis, the decision to publish the manuscript, or the preparation of the manuscript.

## AUTHOR CONTRIBUTIONS

Conceptualization: A.K. and T.W.

Investigation: T.W., A.K., A.S., Y.D., T.K., and T.Mas.

Methodology: T.W., A.K., Y.D., T.K., T.Mas., H.I., M.K., and T.Mor.

Formal analysis: T.W. and A.K.

Resources: H.I., M.K., T.Mas., Y.D., T.K., and T.Mor.

Supervision: A.K.

Writing – original draft: T.W. and A.K.

Writing – review & editing: all authors.

## DATA AVAILABILITY

The RNA-seq data generated in this study have been deposited in the DDBJ Sequence Read Archive under accession numbers DRR751406–DRR751414. The raw proteomics data have been deposited in the jPOST repository under accession number PXD069778. Other data supporting the findings of this study are available within the article and its supplementary material.

## SUPPLEMENTARY DATA

Supplemental material is available for this article: TABLES S1–S5 and FIGS. S1–S13.

## REFERENCES

1. Daly MJ. 2009. A new perspective on radiation resistance based on Deinococcus radiodurans. Nature Reviews Microbiology 2009 7:3 7:237–245.

2. Musilova M, Wright G, Ward JM, Dartnell LR. 2015. Isolation of Radiation-Resistant Bacteria from Mars Analog Antarctic Dry Valleys by Preselection, and the Correlation between Radiation and Desiccation Resistance. Astrobiology 15:1076.

3. Karley D, Shukla SK, Rao TS. 2022. Microbiological assessment of spent nuclear fuel pools: An in-perspective review. J Environ Chem Eng 10:108050.

4. Warashina T, Sato A, Hinai H, Shaikhutdinov N, Shagimardanova E, Mori H, Tamaki S, Saito M, Sanada Y, Sasaki Y, Shimada K, Dotsuta Y, Kitagaki T, Maruyama S, Gusev O, Narumi I, Kurokawa K, Morita T, Ebisuzaki T, Nishimura A, Koma Y, Kanai A. 2024. Microbiome analysis of the restricted bacteria in radioactive element-containing water at the Fukushima Daiichi Nuclear Power Station. Appl Environ Microbiol 90.

5. Naruki M, Watanabe A, Warashina T, Morita T, Arakawa K. 2024. Complete genome sequence of Limnobacter thiooxidans CS-K2 T , isolated from freshwater lake sediments in Bavaria, Germany . Microbiol Resour Announc 13.

6. Hayoun K, Pible O, Petit P, Allain F, Jouffret V, Culotta K, Rivasseau C, Armengaud J, Alpha-Bazin B. 2020. Proteotyping Environmental Microorganisms by Phylopeptidomics: Case Study Screening Water from a Radioactive Material Storage Pool. Microorganisms 2020, Vol 8, Page 1525 8:1525.

7. Spring S, Kämpfer P, Schleifer KH. 2001. Limnobacter thiooxidans gen. nov., sp. nov., a novel thiosulfate-oxidizing bacterium isolated from freshwater lake sediment. Int J Syst Evol Microbiol 51:1463–1470.

8. Friedberg EC. 2003. DNA damage and repair. Nature 2003 421:6921 421:436–440.

9. White O, Eisen JA, Heidelberg JF, Hickey EK, Peterson JD, Dodson RJ, Haft DH, Gwinn ML, Nelson WC, Richardson DL, Moffat KS, Qin H, Jiang L, Pamphile W, Crosby M, Shen M, Vamathevan JJ, Lam P, McDonald L, Utterback T, Zalewski C, Makarova KS, Aravind L, Daly MJ, Minton KW, Fleischmann RD, Ketchum KA, Nelson KE, Salzberg S, Smith HO, Venter JC, Fraser CM. 1999. Genome sequence of the radioresistant bacterium Deinococcus radiodurans R1. Science (1979) 286:1571–1577.

10. Zivanovic Y, Armengaud J, Lagorce A, Leplat C, Guérin P, Dutertre M, Anthouard V, Forterre P, Wincker P, Confalonieri F. 2009. Genome analysis and genome-wide proteomics of Thermococcus gammatolerans, the most radioresistant organism known amongst the Archaea. Genome Biol 10:1–23.

11. Slade D, Radman M. 2011. Oxidative stress resistance in Deinococcus radiodurans. Microbiol Mol Biol Rev 75:133–191.

12. Greenacre M, Groenen PJF, Hastie T, D’Enza AI, Markos A, Tuzhilina E. 2022. Principal component analysis. Nature Reviews Methods Primers 2022 2:1 2:100-.

13. Cox MM. 2007. Regulation of bacterial RecA protein function. Crit Rev Biochem Mol Biol 42:41–63.

14. Drees JC, Lusetti SL, Cox MM. 2004. Inhibition of RecA Protein by the Escherichia coli RecX Protein: MODULATION BY THE RecA C TERMINUS AND FILAMENT FUNCTIONAL STATE. Journal of Biological Chemistry 279:52991–52997.

15. Tatusov RL, Galperin MY, Natale DA, Koonin E V. 2000. The COG database: a tool for genome-scale analysis of protein functions and evolution. Nucleic Acids Res 28:33.

16. West SC, Connolly B. 1992. Biological roles of the Escherichia coli RuvA, RuvB and RuvC proteins revealed. Mol Microbiol 6:2755–2759.

17. Nikaido H. 2003. Molecular basis of bacterial outer membrane permeability revisited. Microbiol Mol Biol Rev 67:593–656.

18. Mah TFC, O’Toole GA. 2001. Mechanisms of biofilm resistance to antimicrobial agents. Trends Microbiol 9:34–39.

19. Liu Y, Beyer A, Aebersold R. 2016. On the Dependency of Cellular Protein Levels on mRNA Abundance. Cell 165:535–550.

20. Esquilin-Lebron K, Dubrac S, Barras F, Boyd JM. 2021. Bacterial Approaches for Assembling Iron-Sulfur Proteins. mBio 12.

21. Varadi M, Anyango S, Deshpande M, Nair S, Natassia C, Yordanova G, Yuan D, Stroe O, Wood G, Laydon A, Zídek A, Green T, Tunyasuvunakool K, Petersen S, Jumper J, Clancy E, Green R, Vora A, Lutfi M, Figurnov M, Cowie A, Hobbs N, Kohli P, Kleywegt G, Birney E, Hassabis D, Velankar S. 2022. AlphaFold Protein Structure Database: massively expanding the structural coverage of protein-sequence space with high-accuracy models. Nucleic Acids Res 50:D439–D444.

22. Teufel F, Almagro Armenteros JJ, Johansen AR, Gíslason MH, Pihl SI, Tsirigos KD, Winther O, Brunak S, von Heijne G, Nielsen H. 2022. SignalP 6.0 predicts all five types of signal peptides using protein language models. Nature Biotechnology 2022 40:7 40:1023–1025.

23. Okuda S, Tokuda H. 2011. Lipoprotein sorting in bacteria. Annu Rev Microbiol 65:239–259.

24. Singleton MR, Dillingham MS, Wigley DB. 2007. Structure and mechanism of helicases and nucleic acid translocases. Annu Rev Biochem 76:23–50.

25. Leipe DD, Wolf YI, Koonin E V., Aravind L. 2002. Classification and evolution of P-loop GTPases and related ATPases. J Mol Biol 317:41–72.

26. Walker JE, Saraste M, Runswick MJ, Gay NJ. 1982. Distantly related sequences in the alpha- and beta-subunits of ATP synthase, myosin, kinases and other ATP-requiring enzymes and a common nucleotide binding fold. EMBO J 1:945–951.

27. Dillingham MS, Kowalczykowski SC. 2008. RecBCD enzyme and the repair of double-stranded DNA breaks. Microbiol Mol Biol Rev 72:642–671.

28. Whittaker CA, Hynes RO. 2002. Distribution and evolution of von Willebrand/integrin A domains: Widely dispersed domains with roles in cell adhesion and elsewhere. Mol Biol Cell 13:3369–3387.

29. Mayer MP, Bukau B. 2005. Hsp70 chaperones: Cellular functions and molecular mechanism. Cell Mol Life Sci 62:670.

30. O’Leary NA, Wright MW, Brister JR, Ciufo S, Haddad D, McVeigh R, Rajput B, Robbertse B, Smith-White B, Ako-Adjei D, Astashyn A, Badretdin A, Bao Y, Blinkova O, Brover V, Chetvernin V, Choi J, Cox E, Ermolaeva O, Farrell CM, Goldfarb T, Gupta T, Haft D, Hatcher E, Hlavina W, Joardar VS, Kodali VK, Li W, Maglott D, Masterson P, McGarvey KM, Murphy MR, O’Neill K, Pujar S, Rangwala SH, Rausch D, Riddick LD, Schoch C, Shkeda A, Storz SS, Sun H, Thibaud-Nissen F, Tolstoy I, Tully RE, Vatsan AR, Wallin C, Webb D, Wu W, Landrum MJ, Kimchi A, Tatusova T, DiCuccio M, Kitts P, Murphy TD, Pruitt KD. 2016. Reference sequence (RefSeq) database at NCBI: current status, taxonomic expansion, and functional annotation. Nucleic Acids Res 44:D733–D745.

31. Thomas CM, Nielsen KM. 2005. Mechanisms of, and Barriers to, Horizontal Gene Transfer between Bacteria. Nature Reviews Microbiology 2005 3:9 3:711–721.

32. Lu H, Sato Y, Fujimura R, Nishizawa T, Kamijo T, Ohta H. 2011. Limnobacter litoralis sp. nov., a thiosulfate-oxidizing, heterotrophic bacterium isolated from a volcanic deposit, and emended description of the genus Limnobacter. Int J Syst Evol Microbiol 61:404–407.

33. Chen Y, Feng X, He Y, Wang F. 2016. Genome analysis of a Limnobacter sp. identified in an anaerobic methane-consuming cell consortium. Front Mar Sci 3:211715.

34. Bushnell B, Rood J, Singer E. 2017. BBMerge – Accurate paired shotgun read merging via overlap. PLoS One 12:e0185056.

35. Pertea G, Pertea, Geo, Pertea, Geo. 2015.fqtrim: v0.9.4 release. zndo 10.5281/ZENODO.20552.

36. Langdon WB. 2015. Performance of genetic programming optimised Bowtie2 on genome comparison and analytic testing (GCAT) benchmarks. BioData Min 8:1–7.

37. Liao Y, Smyth GK, Shi W. 2014. featureCounts: an efficient general purpose program for assigning sequence reads to genomic features. Bioinformatics 30:923–930.

38. Love MI, Huber W, Anders S. 2014. Moderated estimation of fold change and dispersion for RNA-seq data with DESeq2. Genome Biol 15:1–21.

39. Cantalapiedra CP, Hern̗andez-Plaza A, Letunic I, Bork P, Huerta-Cepas J. 2021. eggNOG-mapper v2: Functional Annotation, Orthology Assignments, and Domain Prediction at the Metagenomic Scale. Mol Biol Evol 38:5825–5829.

40. Masuda T, Saito N, Tomita M, Ishihama Y. 2009. Unbiased Quantitation of Escherichia coli Membrane Proteome Using Phase Transfer Surfactants. Molecular & Cellular Proteomics 8:2770–2777.

41. Masuda T, Tomita M, Ishihama Y. 2008. Phase Transfer Surfactant-Aided Trypsin Digestion for Membrane Proteome Analysis. J Proteome Res 7:731–740.

42. Rappsilber J, Mann M, Ishihama Y. 2007. Protocol for micro-purification, enrichment, pre-fractionation and storage of peptides for proteomics using StageTips. Nature Protocols 2007 2:8 2:1896–1906.

43. Virtanen P, Gommers R, Oliphant TE, Haberland M, Reddy T, Cournapeau D, Burovski E, Peterson P, Weckesser W, Bright J, van der Walt SJ, Brett M, Wilson J, Millman KJ, Mayorov N, Nelson ARJ, Jones E, Kern R, Larson E, Carey CJ, Polat İ, Feng Y, Moore EW, VanderPlas J, Laxalde D, Perktold J, Cimrman R, Henriksen I, Quintero EA, Harris CR, Archibald AM, Ribeiro AH, Pedregosa F, van Mulbregt P, Vijaykumar A, Bardelli A Pietro, Rothberg A, Hilboll A, Kloeckner A, Scopatz A, Lee A, Rokem A, Woods CN, Fulton C, Masson C, Häggström C, Fitzgerald C, Nicholson DA, Hagen DR, Pasechnik D V., Olivetti E, Martin E, Wieser E, Silva F, Lenders F, Wilhelm F, Young G, Price GA, Ingold GL, Allen GE, Lee GR, Audren H, Probst I, Dietrich JP, Silterra J, Webber JT, Slavič J, Nothman J, Buchner J, Kulick J, Schönberger JL, de Miranda Cardoso JV, Reimer J, Harrington J, Rodríguez JLC, Nunez-Iglesias J, Kuczynski J, Tritz K, Thoma M, Newville M, Kümmerer M, Bolingbroke M, Tartre M, Pak M, Smith NJ, Nowaczyk N, Shebanov N, Pavlyk O, Brodtkorb PA, Lee P, McGibbon RT, Feldbauer R, Lewis S, Tygier S, Sievert S, Vigna S, Peterson S, More S, Pudlik T, Oshima T, Pingel TJ, Robitaille TP, Spura T, Jones TR, Cera T, Leslie T, Zito T, Krauss T, Upadhyay U, Halchenko YO, Vázquez-Baeza Y. 2020. SciPy 1.0: fundamental algorithms for scientific computing in Python. Nature Method 17:261–272.

44. Wickham H. 2016. ggplot2: Elegant Graphics for Data Analysis. Springer-Verlag New York. https://ggplot2.tidyverse.org.

45. Altschul SF, Madden TL, Schäffer AA, Zhang J, Zhang Z, Miller W, Lipman DJ. 1997. Gapped BLAST and PSI-BLAST: a new generation of protein database search programs. Nucleic Acids Res 25:3389–3402.

46. Nguyen VD, Nguyen TH, Tayeen ASM, Laughinghouse HD, Sánchez-Reyes LL, Wiggins J, Pontelli E, Mozzherin D, O’Meara B, Stoltzfus A. 2020. Phylotastic: Improving Access to Tree-of-Life Knowledge With Flexible, on-the-Fly Delivery of Trees. Evol Bioinform Online 16:1176934319899384.

47. Letunic I, Bork P. 2019. Interactive Tree Of Life (iTOL) v4: recent updates and new developments. Nucleic Acids Res 47:W256–W259.

48. Lee MD. 2019. GToTree: a user-friendly workflow for phylogenomics. Bioinformatics 35:4162–4164.

49. Sahl JW, Gregory Caporaso J, Rasko DA, Keim P. 2014. The large-scale blast score ratio (LS-BSR) pipeline: A method to rapidly compare genetic content between bacterial genomes. PeerJ 2014:e332.

50. Jones P, Binns D, Chang HY, Fraser M, Li W, McAnulla C, McWilliam H, Maslen J, Mitchell A, Nuka G, Pesseat S, Quinn AF, Sangrador-Vegas A, Scheremetjew M, Yong SY, Lopez R, Hunter S. 2014. InterProScan 5: genome-scale protein function classification. Bioinformatics 30:1236–1240.

51. Katoh K, Standley DM. 2013. MAFFT Multiple Sequence Alignment Software Version 7: Improvements in Performance and Usability. Mol Biol Evol 30:772–780.

52. Thompson JD, Gibson TJ, Plewniak F, Jeanmougin F, Higgins DG. 1997. The CLUSTAL_X Windows Interface: Flexible Strategies for Multiple Sequence Alignment Aided by Quality Analysis Tools. Nucleic Acids Res 25:4876–4882.

53. Mirdita M, Schütze K, Moriwaki Y, Heo L, Ovchinnikov S, Steinegger M. 2022. ColabFold: making protein folding accessible to all. Nat Methods 19:679–682.

54. van Kempen M, Kim SS, Tumescheit C, Mirdita M, Lee J, Gilchrist CLM, Söding J, Steineggeri M. 2024. Fast and accurate protein structure search with Foldseek. Nat Biotechnol 42:243–246.

55. Pettersen EF, Goddard TD, Huang CC, Meng EC, Couch GS, Croll TI, Morris JH, Ferrin TE. 2020. UCSF ChimeraX: Structure visualization for researchers, educators, and developers. Protein Sci 30:70.

56. Davison J, Heusterspreute M, Chevalier N, Brunei F. 1987. A “phase-shift” fusion system for the regulation of foreign gene expression by lambda repressor in gram-negative bacteria. Gene 60:227–235.

57. Ehrlich SD. 1977. Replication and expression of plasmids from Staphylococcus aureus in Bacillus subtilis. Proc Natl Acad Sci U S A 74:1680.

58. Piffaretti JC, Arini A, Frey J. 1988. pUB307 mobilizes resistance plasmids from Escherichia coli into Neisseria gonorrhoeae. Mol Gen Genet 212:215–218.

59. Datsenko KA, Wanner BL. 2000. One-step inactivation of chromosomal genes in Escherichia coli K-12 using PCR products. Proc Natl Acad Sci U S A 97:6640–6645.

60. Mirdita M, Schütze K, Moriwaki Y, Heo L, Ovchinnikov S, Steinegger M. 2022. ColabFold: making protein folding accessible to all. Nature Methods 2022 19:6 19:679–682.

61. Valdar WSJ. 2002. Scoring residue conservation. Proteins: Structure, Function and Genetics 48:227–241.

